# Discerning role of a functional arsenic resistance cassette in evolution and adaptation of a rice pathogen

**DOI:** 10.1101/2020.12.16.422644

**Authors:** Amandeep Kaur, Rekha Rana, Tanu Saroha, Prabhu B. Patil

## Abstract

Arsenic (As) is highly toxic element to all forms of life and is a major environmental contaminant. Understanding acquisition, detoxification, and adaptation mechanisms in bacteria that are associated with host in arsenic-rich conditions can provide novel insights into dynamics of host-microbe-microenvironment interactions. In the present study, we have investigated an arsenic resistance mechanism acquired during the evolution of a particular lineage in the population of *Xanthomonas oryzae* pv. *oryzae* (*Xoo)*, which is a serious plant pathogen infecting rice. Our study revealed the horizontal acquisition of a novel chromosomal 12kb *ars* cassette in *Xoo* IXO1088 that confers high resistance to arsenate/arsenite. The *ars* cassette comprises several genes that constitute an operon induced in the presence of arsenate/arsenite. Transfer of cloned *ars* cassette to *Xoo* BXO512 lacking it confers arsenic resistance phenotype. Further, the transcriptional response of *Xoo* IXO1088 under arsenate/arsenite exposure was analyzed using RNA sequencing. Arsenic detoxification and efflux, oxidative stress, iron acquisition/storage, and damage repair are the main cellular responses to arsenic exposure. Our investigation has provided novel insights in to how a pathogenic bacterium is coping with arsenic-rich unique micro-environments like seen in rice growing in submerged water conditions.

**Impact statement:** Arsenic accumulation in rice is a serious and unique agronomic issue. Arsenic contaminated groundwater used for irrigation purposes is adding to the accumulation of arsenic in rice. Submerged conditions in the paddy fields further induce the prevalence of toxic inorganic arsenic species in the environment. Our genomics and transcriptomics-based study reveals how a rice pathogen is coping with the lethal concentrations of arsenic by acquiring a novel resistance cassette during diversification into lineages. Acquisition of such detoxification mechanisms can provide a selective advantage to the bacterial population in avoiding toxicity or enhancing virulence and to their on-going evolutionary events. While there are numerous studies on plant-pathogen-environment interactions, our study highlights the importance of systematic studies on the role of unique micro-environmental conditions on the evolution of host-adapted pathogens/microbes.

## Introduction

Arsenic is a highly toxic heavy metal that occurs in the natural environment either through natural processes (volcanic eruption, mineral activities) or by anthropogenic activities (use of pesticides, fertilizers, industrial activities, and agricultural practices) (Zhu et al., 2014) (Naidu and Bhattacharya, 2009). Arsenic exists in different forms in the environment (organic and inorganic), but arsenate [As(V)] and arsenite [As(III)] are two predominant inorganic forms that are toxic to the cells. Arsenite (III) is more toxic than arsenate (V) and mostly predominates under anoxic and reduced environment, whereas arsenate predominates in soil and oxygenated surface water (Hughes, 2002). Accordingly, bacteria have evolved different mechanisms for As-detoxification including As-reduction, As-oxidation, extrusion of As (III) out of cells mediated by efflux transporters, As-methylation, etc. (Yan et al., 2019) (Slyemi and Bonnefoy, 2012) (Yang et al., 2012) (Oremland and Stolz, 2003). These resistance mechanisms in the bacterial population are encoded by the presence of *ars* operons that include *arsRBC*, *arsRABC*, and *arsRDABC* (Ben Fekih et al., 2018) (Yang et al., 2012). In some cases, *ars* genes also exist in a single copy. These operons are either chromosomally encoded or plasmid-encoded and are usually acquired through horizontal gene transfer events (Ben Fekih et al., 2018). The major genes in the operon include *arsR* gene that encodes transcriptional regulator, *arsC* encodes arsenate reductase that transforms arsenate to arsenite, *arsB* encodes a membrane-bound arsenite carrier protein ArsB that efflux arsenite out of the cell, and *arsA* codes for ATPase subunit that forms a complex with ArsB to form arsenite efflux pump (Yan et al., 2019) (Yang et al., 2012). The complex operons also include *arsD*, which encodes for metallo-chaperone to deliver arsenite to ArsA. Other than these, recently characterized *ars* genes include *arsH*, *arsM, arsP*, and *arsI* which broaden resistance towards organic arsenicals (Li et al., 2016) (Yang and Rosen, 2016). The *arsH* gene encodes for an organoarsenical oxidase enzyme that confers resistance to methyl As(III) (Chen et al., 2015). The *arsM* gene encoding ArsM (arsenite S-adenosylmethionine methyltransferase) methylates arsenite to volatile trimethyl arsenite (Qin et al., 2006).

Nowadays, Arsenic contamination from agricultural sources is a growing concern. In countries like India, Bangladesh, Argentina, and China, As-contaminated groundwater used for irrigation adds to the accumulation of As in soil and uptake of As by crop plants (Rahaman et al., 2013) (Ahmad et al., 2018) (Meharg and Rahman, 2003) (Alam and Sattar, 2000). Accumulation of As in rice has become a great disaster as rice is a staple food for half of the world population (Zhu et al., 2008) (Mondal and Polya, 2008) (Meharg et al., 2009) (Ma et al., 2008). Unlike in other crops, the anaerobic conditions prevailing in the paddy field make the environment suitable for As accumulation. Usually, plant species have successfully adapted different mechanisms to cope with toxic metal’s toxic effects, such as detoxification, elimination, and heavy metals accumulation. Studies have elucidated the mechanisms of As uptake and its translocation to different parts of rice plants, including roots, shoots, grains, and seeds (Ma et al., 2008) (Wu et al., 2011). Besides affecting plants’ systems, arsenic present in the environment can also positively or negatively affect the bacterial population that grows in association with plants (Poschenrieder et al., 2006). Response to environmental stress specifically posed by toxic chemicals allows bacteria to evolve under high selection pressure or acquire resistance against toxic substances (Prabhakaran et al., 2016). Some of the bacteria that co-evolve with plants or exposure to heavy metals have resulted in the acquisition of resistance mechanisms to cope with metal-toxicity (Chen et al., 2016) (Lu et al., 2017). These resistance mechanisms help bacteria to not only prevent metal-toxicity but also provide cross-protection against plant defense responses. Studies have shown that arsenite exposure can induce oxidative stress in bacteria leading to the expression of stress-inducible genes (Andres and Bertin, 2016). Induction of such responses can provide a survival advantage to the bacteria during plant-pathogen interactions. Various antioxidant enzymes that are induced on exposure to arsenite can provide cross-protection against reactive oxygen species (ROS), which are generated by plants as an initial defense response against invading pathogens (Hrimpeng et al., 2006) (Sukchawalit et al., 2005).

In the present study, we demonstrate the arsenic resistance mechanisms in *Xanthomonas oryzae* pv. *oryzae* (*Xoo*) population that causes bacterial blight disease in rice plants and is a serious threat to rice cultivation. As *Xoo* is a pathogen of rice, it is likely that *Xoo* population is exposed to an arsenic environment and is under selection for adaptation to higher arsenic concentration, particularly found in the submerged rice cultivation conditions. Inspection of the genomic sequence of a *Xoo* strain IXO1088 revealed the presence of a 12kb *ars* cassette acquired through horizontal gene transfer events and expanded in a particular lineage of *Xoo* population. The cassette confers high arsenic resistance to *Xoo* strain IXO1088 carrying *ars* cassette, whereas a *Xoo* strain BXO512 lacking this cassette was sensitive to the arsenic ions.

Cloning and transfer of *ars* cassette rescued the sensitive strain BXO512 from the lethal concentration of arsenic. The transcriptional analysis demonstrated that the expression of *ars* operon is induced by arsenate and arsenite and is regulated by *arsR*. Further, to investigate the genome-scale transcriptional response of *Xoo* to arsenic stress, we performed RNA sequencing under both As(V) and As(III) exposure. Our results provided the set of differentially expressed genes under both conditions highlighting the mechanisms employed by *Xoo* in response to arsenic treatment. Overall, our study shows how arsenic resistance mediated by horizontal gene transfer of a novel arsenic resistance cassette shaping the population structure and lineage of a rice pathogen. At the same time, the study; also highlights the importance of the plant-specific microenvironment in host-pathogen interaction.

## Results

### *X. oryzae* pv. *oryzae* IXO1088 harbors a novel arsenic resistance cassette

In the present study, we identified a unique 12,214 bp *ars* cassette in *Xoo* strain IXO1088 genome related to the arsenic resistance **Figure 1(A)**. Genes in the cassette include transcriptional regulator, two *arsC* genes, which encode for arsenate reductase, *arsH* gene that encodes for arsenical resistance protein, arsenic transporter, AAA family ATPase, sulfurtransferase and site-specific integrase. The cassette is flanked by transposases on both sides and a site-specific integrase on one end, indicating its acquisition through horizontal gene transfer events. Then, we looked into the cassette‘s GC content, and the presence of genes with atypical GC content further suggested the acquisition of this cassette from other organisms **Supplementary Table 1**. A more detailed analysis of the cassette revealed a putative promoter region recognized by a sigma 70 transcription factor in the 5’ upstream of the helix-turn-helix transcriptional regulator **Figure 1(B)**. The other *ars* genes in the cassette *arsC, arsH,* gene encoding arsenic transporter, do not contain promoter regions. This gene organization suggested that the *ars* cassette constitutes an operon, whose expression is regulated by helix-turn-helix transcriptional regulator. The transcriptional regulator present in the cassette showed homology to the ArsR/Smt family of transcriptional regulators. Sequence alignment of the regulator with ArsR family proteins from other bacteria indicates conservation of helix-turn-helix domain **Figure 1(C)**. This suggested that the transcriptional regulator in the cassette belongs to ArsR family regulators and its regulation by arsenic ions.

**Figure 1:**
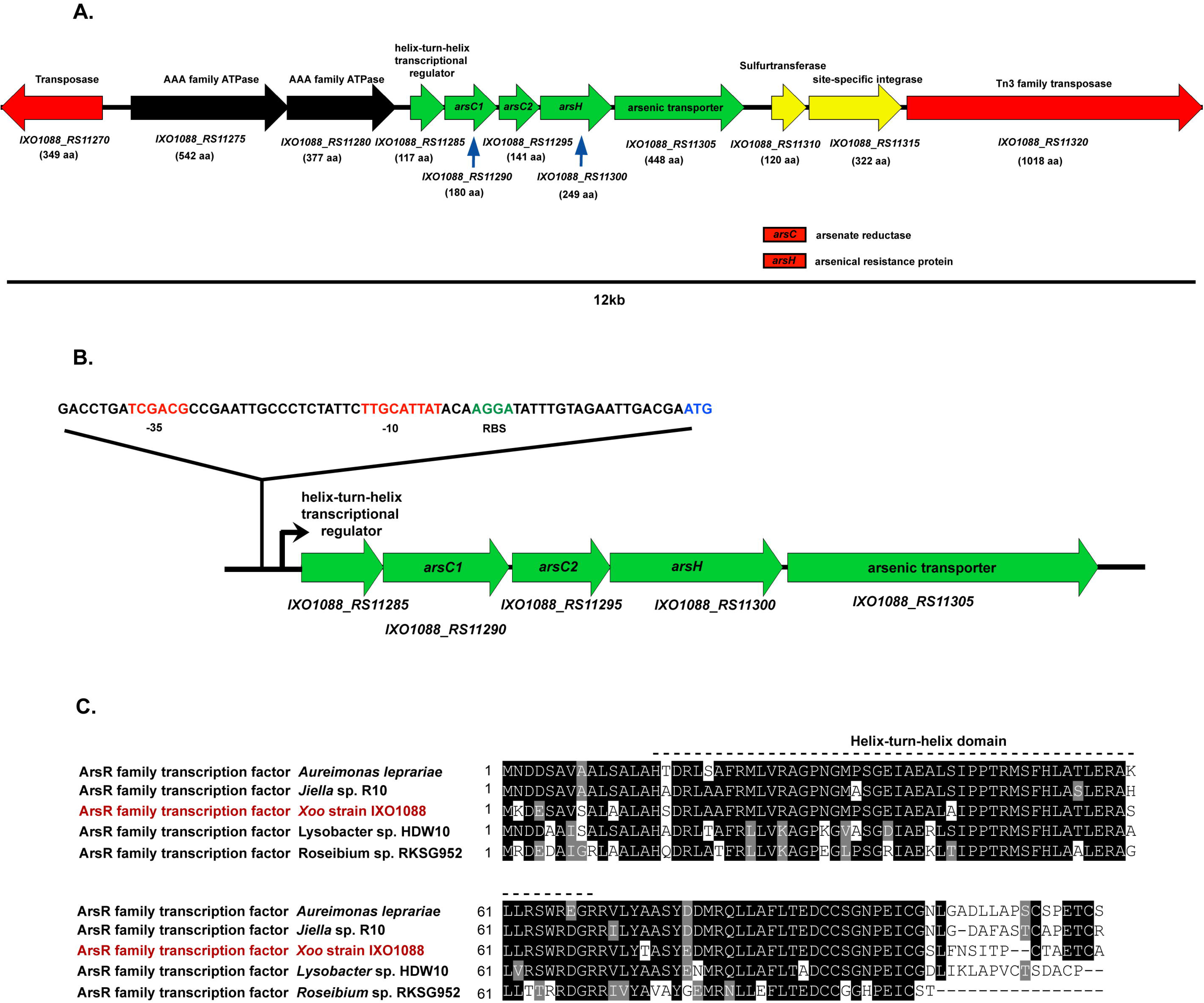
Genetic organization of *ars* operon present in *Xoo* strain IXO1088. **(A)** The arrows represent genes present in the cassette and the direction of the transcription. Below arrows, locus tags are given for each gene along with the size (number of amino-acid residues). **(B)** The putative promoter sequence upstream of *arsR* gene with the predicted conserved regions −35 and −10 highlighted in red color; RBS (ribosome binding site) in green and initiation codon in blue. **(C)** Multiple sequence alignment of ArsR family regulator of IXO1088 with the ArsR family transcription factor homologs from other bacterial species. The dashed line represent helix-turn-helix domain.The black background indicates identical amino acid residues.

### Phylogenetic relatedness of arsenate reductase and arsenic transporter genes

To identify the related neighbors of arsenate reductase genes (*IXO1088_11290* and *IXO1088_11295*) and arsenite transporter (*IXO1088_11305*), we performed the phylogenetic analysis of these proteins with the *arsC* and *arsB* proteins form various arsenic-resistant microorganisms, respectively. The phylogenetic tree of ArsC protein sequences revealed that the two ArsC proteins encoded in the cassette fall into two groups, one ArsC1 protein (IXO1088_11290) fall in the ArsC arsenate reductases of thioredoxin (Trx) family whereas, the other ArsC2 protein (IXO1088_11295) grouped with the arsenate reductases of glutaredoxin (Grx) family **Figure 2(A).** Usually, the primitive *arsRBC* operon comprises ArsB arsenite efflux pump that extrudes arsenite from the cells. Later on, it was found that *Bacillus subtilis* contained a novel arsenite transporter that showed homology to Acr3 protein from yeast *Saccharomyces cerevisiae,* which confers arsenic resistance. The *acr3* gene was subsequently found in many microorganisms such as *R. palustris, C. jejuni, H. arsenioxydans, Microbacterium* sp. etc. (Ben Fekih et al., 2018). The phylogenetic analysis of arsenic transporter (*IXO1088_11305*) present in IXO1088 cassette revealed that it is closely related to ArsB family transporters and all Arc3 transporters fall in other clade **Figure 2(B)**.

**Figure 2:**
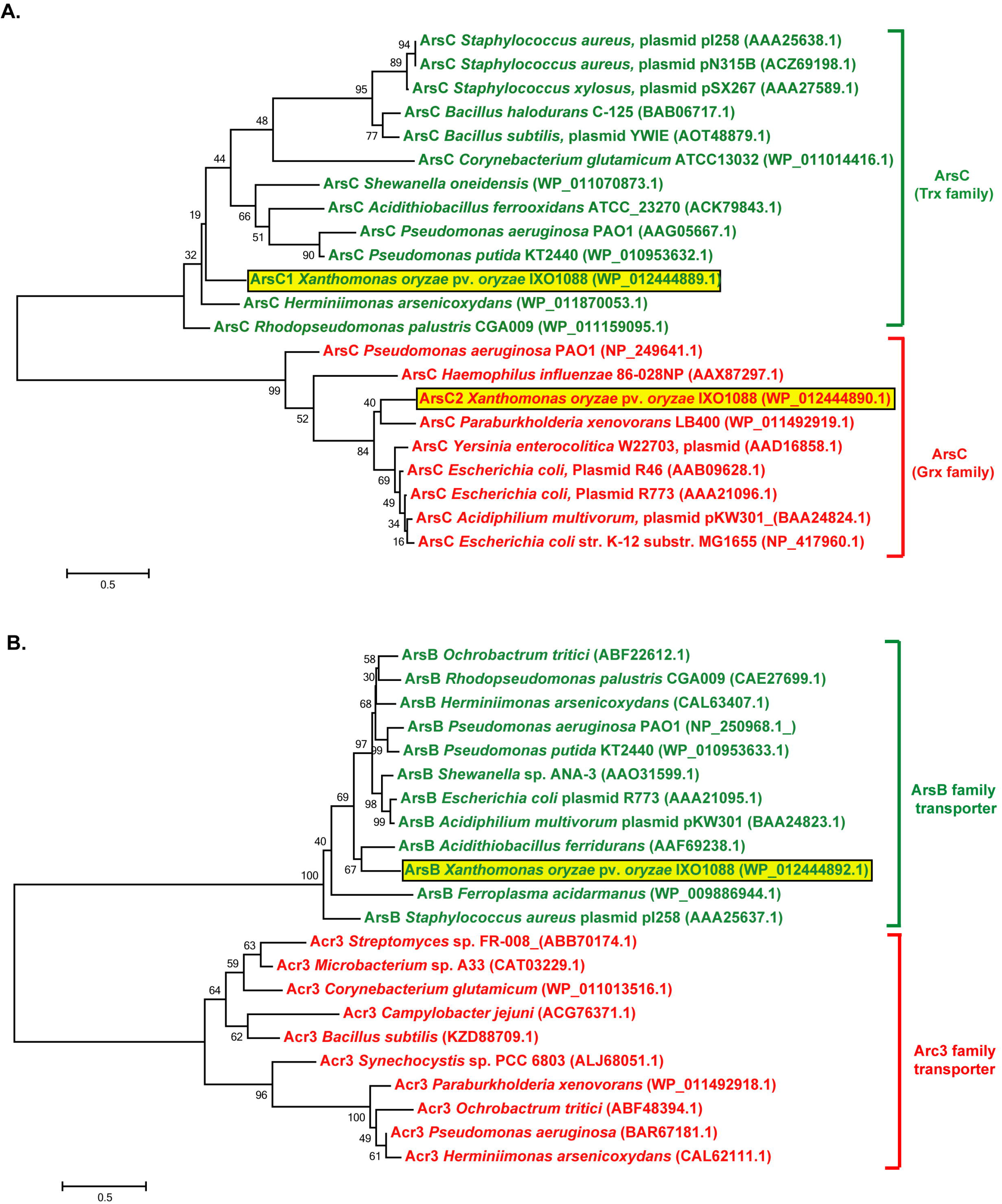
Phylogenetic analysis of arsenate reductase (ArsC) and arsenic transporter proteins encoded in the *ars* cassette of *Xoo* strain IXO1088 strain. Phylogenetic tree of **(A)** ArsC protein and **(B)** arsenic transporter protein from IXO1088 with other bacterial species constructed using MEGA6 software. The *Xoo* IXO1088 ArsC and ArsB proteins are highlighted in a yellow box, and sequence accession numbers for all are given in parentheses. Bootstrap values are shown at each branch point.

### Distribution of *ars* cassette in *X. oryzae* pv. *oryzae* population

Core genome phylogenetic analysis of 413 *Xoo* genomes available in NCBI from different geographical regions was performed, and then, we looked for the presence of *ars* cassette in all the genomes. Phylogenetic analysis revealed six distinct lineages of *Xoo* population. Interestingly, we observed that this cassette was present in a particular lineage comprising 55 isolates **Figure 3**. The lineage predominantly constitutes strains from Asia (53), including strains from India (22), China (21), Philippines (9), and Nepal (1). One strain from Colombia and Bolivia each was also present. Further, we observed that in India, the strains carrying *ars* cluster were isolated from areas where arsenic contamination in groundwater or irrigation water is a problem **Supplementary Figure 1**.

**Figure 3:**
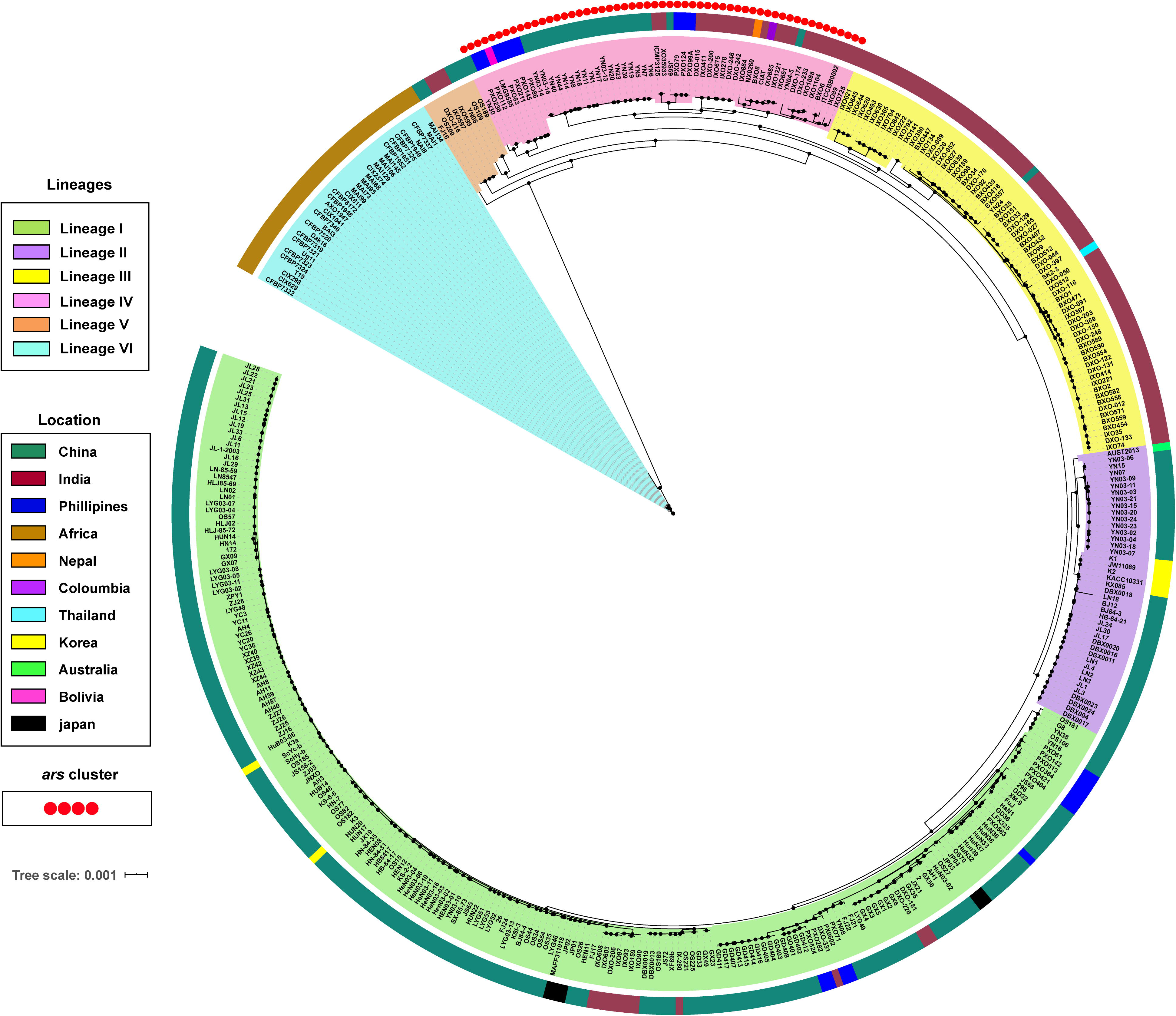
Distribution of *ars* cassette among *Xoo* population. The core genome tree was constructed using PhyML and visualized using iTOL software. Background colors of nodes represent different lineages. The innermost circle represents different countries from where strains were isolated, and the outermost circle (red circles) refers to strains in which the *ars* cassette is present.

### Regulation and functional characterization of arsenic resistance gene cassette present in *X. oryzae* pv. *oryzae* IXO1088

Arsenic sensitivity assay showed that *Xoo* IXO1088 could tolerate significantly higher concentrations of arsenate and arsenite compared to another *Xoo* BXO512 that lacks *ars* cassette. Significant growth inhibition of IXO1088 strain to arsenic ions was only observed above 12mM arsenate and 0.6 mM arsenite **Figure 4(A) and 4(B)**. Compared to IXO1088 strain, BXO512 strain could not grow even in the presence of 0.1 mM arsenite **Figure 4(B)**. However, BXO512 strain tolerated a slight concentration of arsenate, but significant inhibition was observed even at 0.5 mM concentration of arsenate **Figure 4(A)**. A single copy of arsenate reductase gene in the BXO512 genome could contribute to the slight tolerance to arsenate. We found that a single copy of arsenate reductase gene is present in all the *Xoo* genomes that can provide resistance against that basal concentration of arsenate. To check whether arsenic resistance phenotype is directly linked with the *ars* cassette, we cloned the complete *ars* gene cluster in pUFR034 cosmid vector, transferred recombinant plasmid (pUFR034+*ars*) to the BXO512 strain, and performed arsenic sensitivity assay. As expected, BXO512 strain with this cassette was resistant to both arsenate and arsenite as compared to the strain with only empty vector (pUFR034) **Figure 4(C) and 4(D)**. The plasmid presence not only increased arsenate resistance but also conferred an increased level of resistance against arsenite. This data indicated that *Xoo* IXO1088 *ars* cassette is functional when transferred to the other *Xoo* strain.

**Figure 4:**
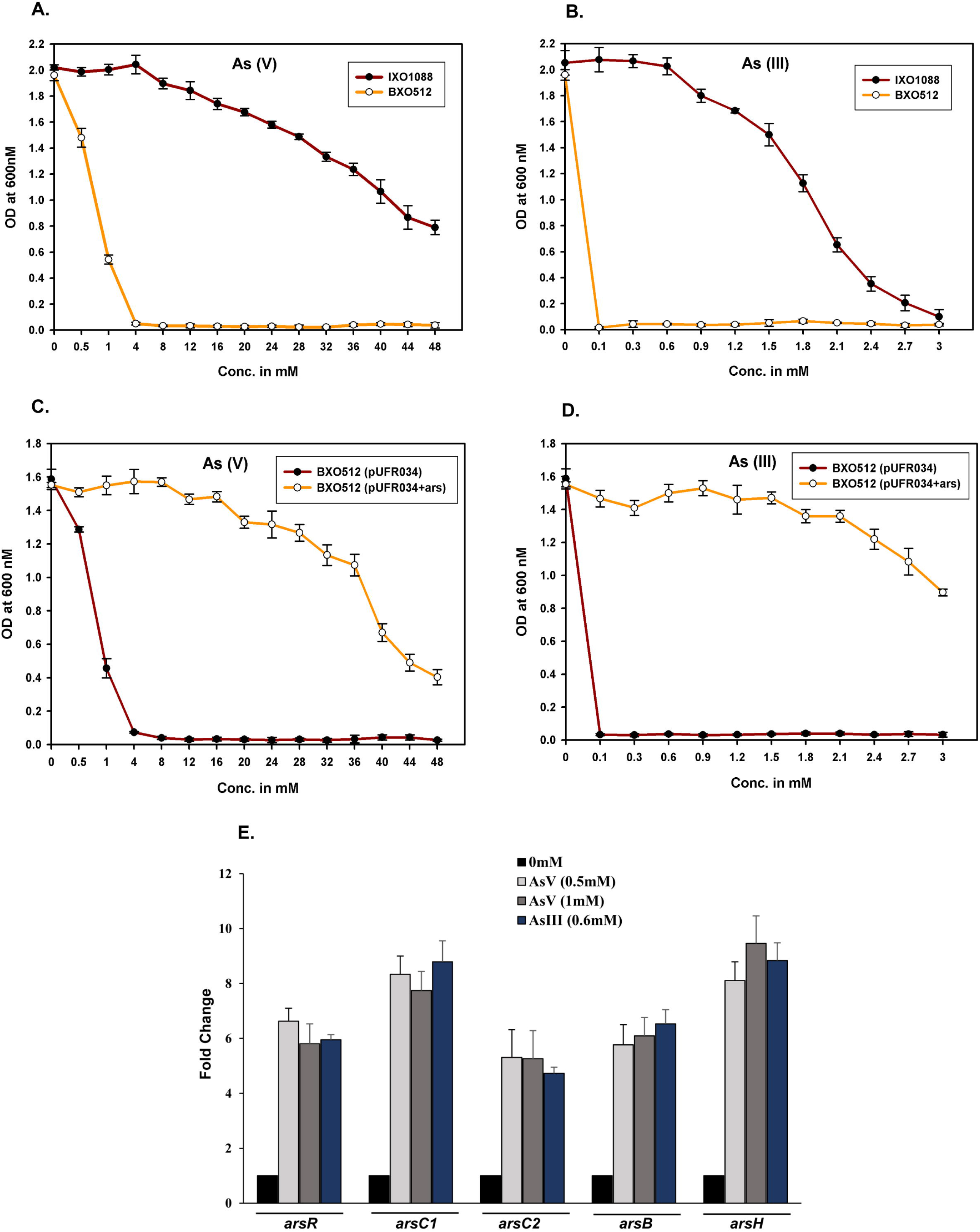
Arsenic sensitivity assay of *Xoo* strain IXO1088 (contains *ars* cassette) and *Xoo* strain BXO512 (does not contains *ars* cassette). Cultures were grown in nutrient broth (NB) with different concentrations of **(A)** sodium arsenate and **(B)** sodium arsenite. The below panel shows susceptibility of *Xoo* strain BXO512 containing *Xoo* IXO1088 *ars* cassette against **(C)** sodium arsenate and **(D)** sodium arsenite, wild type *Xoo* BXO512 (pUFR034) (orange color); *Xoo* BXO512 (pUFR034+ars cassette) (brown color). Data values shown are means with standard deviation from three independent experiments. **(E)** The *ars* operon is induced in the presence of sodium arsenate and sodium arsenite. Transcription of *arsR, arsC1, arsC2, arsB*, and *arsH* genes was determined using qRT-PCR in cells grown in the presence of arsenate (0.5mM and 1mM) and arsenite (0.6mM) during exponential phase. Relative expression was calculated using ΔΔct method. ftsZ was used as an endogenous control. Error bars represent the standard deviation from three independent biological replicates (each performed in three technical replicates).

To determine whether the arsenic ions regulates the expression of *ars* cassette, quantitative reverse transcriptase PCR (qRT-PCR) was used to compare the transcription of the *arsR*, *arsC1*, *arsC2*, *arsH*, and *arsB* genes in *Xoo* IXO1088 grown in the presence and absence of subinhibitory concentrations of arsenate and arsenite (as mentioned in material and methods section). The subinhibitory concentration was decided by growing cells in the presence of different concentrations of arsenate and arsenite by performing growth curve assay **Supplementary Figure 2.** The transcription of *ars* operon was regulated in the presence of both arsenate and arsenite. The expression of *arsR*, *arsC1*, *arsC2*, *arsH*, and *arsB* genes was induced 6 to 8 fold in arsenate and arsenite treated samples compared to the control cells without any treatment **Figure 4(E)**. The data suggested that the presence of arsenic ions induces *ars* operon. The expression level for the other two genes encoding AAA family ATPase remains unchanged (data not shown).

### Transcriptomics of *X. oryzae* pv. *oryzae* IXO1088 under arsenic stress

To study the defense mechanism employed by *Xoo* in response to arsenic treatment, we carried out RNA sequencing at three concentrations two for arsenate (0.5mM and 1mM) and 0.6mM for arsenite, as mentioned in the material and methods section. On average, 30 million reads were obtained for each sample, and more than 98% of the reads were mapped to the reference genome. We further analyzed the differentially expressed genes. Genes with adjusted *p*-value <0.05 were considered significant, and genes with log_2_fold change ≥1 were considered upregulated, and log_2_fold change ≤−1 were considered downregulated **Supplementary Figure 3**. The complete list of DEGs under all three conditions is shown in **Supplementary Table 2.** Further analysis of DEGs revealed that a total of 21 upregulated and 15 downregulated genes. were shared by all three conditions. As As(III) is highly toxic compared to As(V), the number of DEGs in As(III) treated sample was more with 202 upregulated and 204 downregulated genes compared to the As(V) treated samples. Whereas, in 0.5mM As(V) treated sample, 48 up-regulated and 76 down-regulated genes and in the sample treated with 1mM As(V), 45 up-regulated, and 55 down-regulated genes were found **Figure 5(A)**. This shows that 0.5mM and 1mM As(V) concentrations could be a less stressful condition for the cells than As(III).

**Figure 5:**
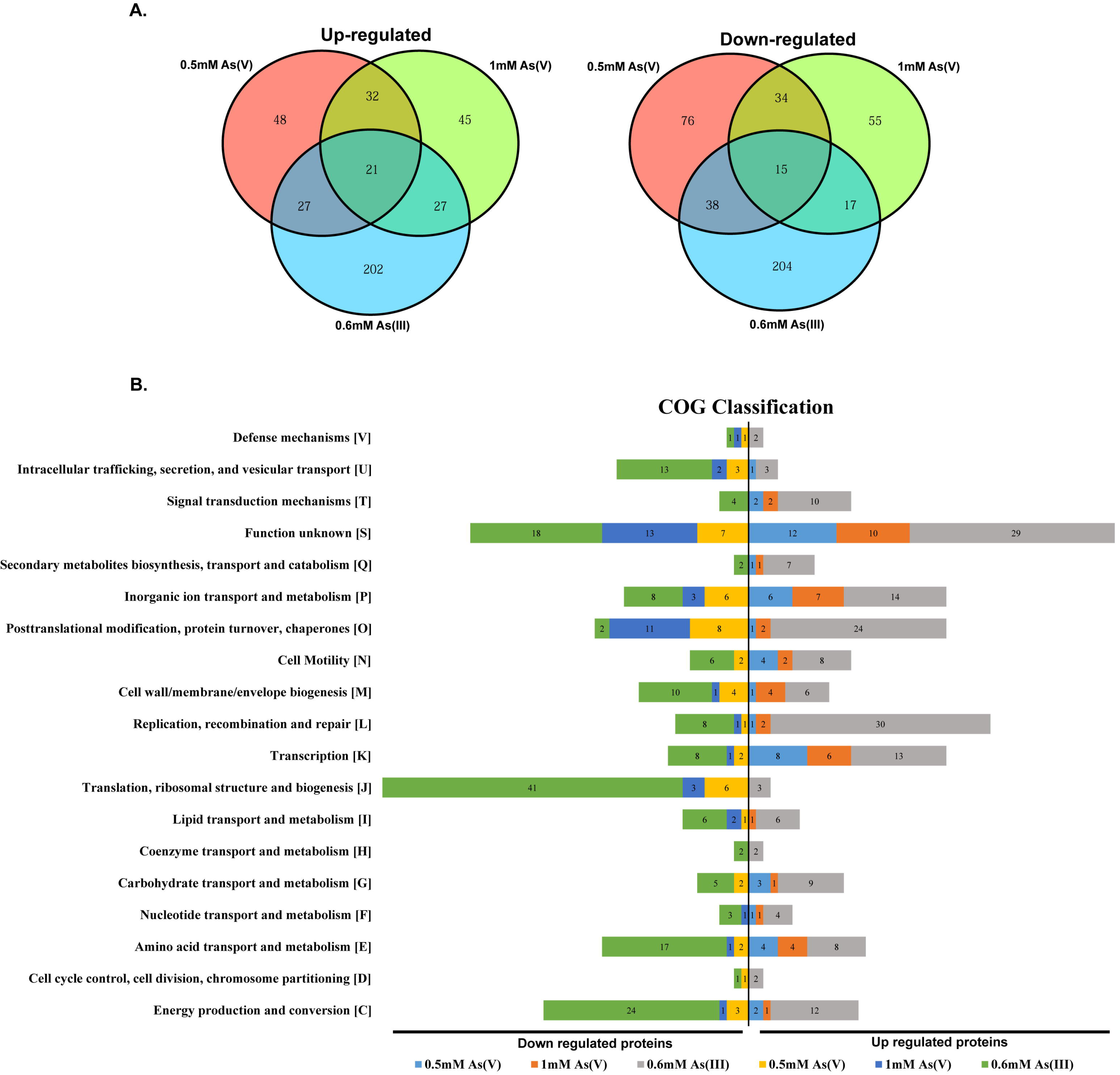
**(A)** Venn diagrams showing DEGs (differentially expressed genes) upregulated or downregulated under arsenate As(V) and arsenite As(III) treatment. **(B)** COG (Cluster of orthologous groups) classification of DEGs upregulated or downregulated under arsenate As(V) and arsenite As(III) treatment. The numbers within the bars represent the number of proteins present in that particular category.

The DEGs were further classified into different COGs (Cluster of orthologous groups). A total of 552 genes were classified into 19 COG categories **Figure 5(B)**. The COG analysis revealed that the maximum number of DEGs fall in the function unknown category (89; 16.1%), followed by other categories including translation, ribosomal structure and biogenesis (53; 9.6%), posttranslational modification, protein turnover, chaperons (48; 8.7%), inorganic ion transport and metabolism (44; 8.0%), energy production and conversion (43; 7.8%), replication, recombination and repair (43; 7.8%), transcription (38; 6.9%), amino acid transport and metabolism (36; 6.5%), cell wall/membrane/envelope biogenesis (26; 4.7%) and intracellular trafficking, secretion, and vesicular transport (22; 4.0%), cell motility (22; 4.0%), carbohydrate transport and metabolism (20; 3.6%), signal transduction mechanisms (18; 3.3%). The other categories with the lowest number include cell motility, nucleotide transport and metabolism, lipid transport and metabolism, Secondary metabolites biosynthesis, transport and catabolism, defense mechanisms, coenzyme transport and metabolism, cell cycle that contributes (50; 9.0%) in total.

### Gene expression pattern common to both arsenate As(V) and arsenite As(III)

Significantly, upregulated genes induced by all three conditions include *ars* operon encoding helix-turn-helix transcriptional regulator (*RS_11285)*, arsenate reductase (*RS_11290)*, arsenate reductase (glutaredoxin) (*RS_11295)*, arsenical resistance protein (*RS_11300)*, arsenic transporter (*RS_11305)* **Figure 6(A)**. The upregulation of the operon in RNA seq experiment was consistent with our induction results. Additionally, other genes that were upregulated, encodes DNA binding protein (*RS_14295)*, abr family transcriptional regulator (*RS_14300)*, MarR transcriptional regulator (*RS_01475)*, (2Fe-2S)-binding protein (*RS_02860)*, tonB dependent siderophore receptor (*RS_06840)*, hemin transporter HemP (*RS_19500)*, BON domain-containing protein (*RS_20250)* **Figure 6(A)**. In contrast, significantly downregulated genes encodes bacterioferritin (*RS_02865* and *RS_09635*), potassium ATPase subunit F (*RS_19820)*, potassium ATPase subunit A (*RS_19815)*, RND-efflux transporter subunits (*RS_15130* and *RS_15135)*, methionine synthase (*RS_10005)* **Figure 6(A)**.

**Figure 6:**
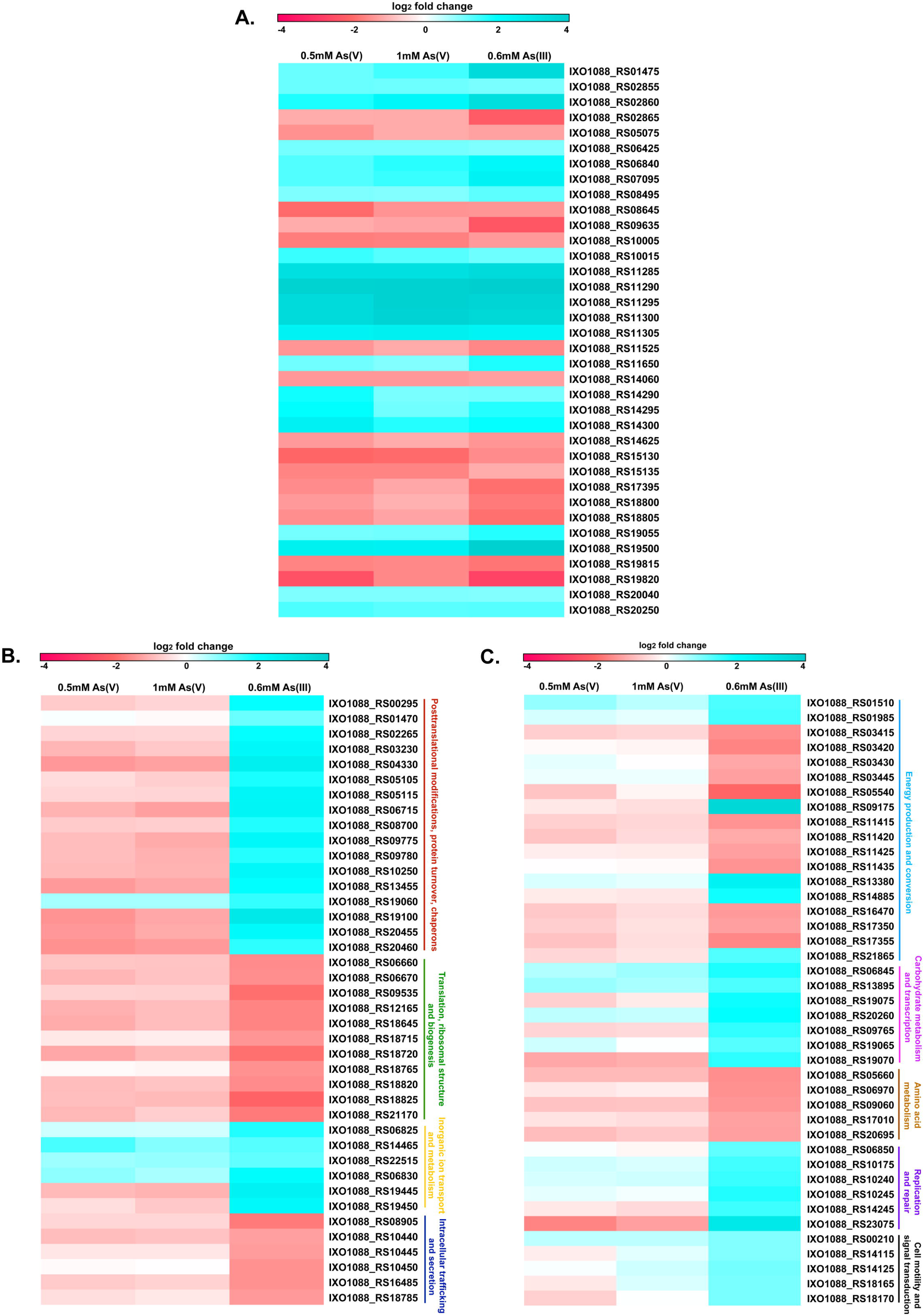
Heat map visualization of up-regulated and down-regulated genes. **(A)** Common differentially expressed genes (up and downregulated) in response to both arsenate and arsenite. **(B)** And **(C)** Heat map visualization of differentially expressed genes in response to arsenite. The scale represents the log_2_ fold change value.

### Gene expression pattern specific to arsenite As(III) exposure

More genes were differentially expressed in response to As(III) treatment, and their expression level was different compared to As(V) treatment **Figure 6(B) and 6(C)**. The genes regulated in response to As(III) treatment categorized into different categories are as below:

#### Oxidative stress proteins

Several genes encoding stress-related proteins and are known to protect against oxidative damage were upregulated in response to As(III) treatment. These genes encodes superoxide dismutase (*RS_13380*), flavin-dependent oxidoreductase (*RS_14885*), serine endopeptidase (*RS_00295*), organic hydroperoxide resistance protein (*RS_01470*), peptide methionine sulfoxide reductase MsrA and MsrB (*RS_19100* and *RS_03230*), alkyl hydroperoxide reductase subunit F (*RS_19060*), oxidative stress transcriptional regulator (*RS_19065*), nucleoside diphosphate kinase regulator (*RS_19070*) and catalase (*RS_22515*) **Figure 6(B)**.

#### Heat shock proteins (Hsps)

Heat shock proteins (Hsps) function as chaperons, are considered as general markers of cell damage, and stress conditions. Hsps facilitate proper folding of newly or already translated proteins and prevents aggregation of inappropriately folded proteins. We observed the upregulation of major Hsps under arsenic stress conditions. Many genes encoding heat-inducible transcriptional repressor HrcA (*RS_09765*), chaperonin GroEL (*RS_02265*), Hsp20/alpha_crystallin_family_protein (*RS_04330*), ATP-dependent Clp endopeptidase proteolytic subunit ClpP (*RS_05105*), endopeptidase La (*RS_05115*), ATP-dependent chaperone ClpB (*RS_06715*), Hsp33 family molecular chaperone HslO (*RS_08700*), molecular chaperone DnaK (*RS_09775*), molecular chaperone DnaJ (*RS_09780*), molecular chaperone htpG (*RS_10250*), protease HtpX (*RS_13455*), ATP-dependent protease ATPase subunit HslU and HslV (*RS_20455* and *RS_20460*) were found to be upregulated **Figure 6(B)**. In addition, we observed downregulation of protein translocase subunits YajC, SecD, F, G and SecE involved in intracellular protein transport (*RS_10440*, *RS_10445*, *RS_10450*, *RS_16485* and *RS_18785*) **Figure 6(B)**.

#### Iron acquisition, storage or production of siderophores

RNA-seq results also revealed significant upregulation of genes related to Fe-S cassettes biogenesis and iron recruitment to alleviate the arsenic stress. Genes related to iron storage were differentially expressed under both AsV and AsIII treated conditions. Fe-S binding protein (*RS_02860*), TonB-dependent siderophore receptor (*RS_06840*), hemin uptake protein HemP (*RS_19500*), MarR family transcriptional regulator (*RS_01475*) were significantly upregulated in all conditions. In contrast, Iron transporter (*RS_06820*), MFS transporter (*RS_06825*), IucA/IucC family siderophore biosynthesis protein (*RS_06830*), AbrB family transcriptional regulator (*RS_14300*), Ferrous iron transport protein (*RS_14465*), iron-sulfur cassette insertion protein ErpA (*RS_21865*) were significantly upregulated in AsIII treated samples **Figure 6(C)**.

#### Recombination and energy metabolism

Interestingly, genes involved in repair and recombination encoding various transposases were found to be upregulated (*RS_06850*), (*RS_10175*), (*RS_10240*), (*RS_10245*), (*RS_14245*), and (*RS_23075*) **Figure 6(C)**. Also, genes involved in electron transfer pathway in oxidative phosphorylation including NADH:Flavin oxidoreductase (*RS_01985*), alkene reductase (*RS_09175*), LLM class Flavin dependent oxidoreductase (*RS_14885*), ATP-binding cassette protein (*RS_19445*), FtsX-like permease family protein (*RS_19450*) were found to be upregulated. In contrast, F0F1 ATP synthase subunits (*RS_03415*, *RS_03420*, *RS_03430* and *RS_03445*), cytochrome ubiquinol oxidase subunits (*RS_06400*, *RS_06405*, *RS_17350*, *RS_17355* and *RS_17360*) were found to be downregulated in response to arsenite **Figure 6(C)**. Other than this, genes involved in translation machinery encoding 50S ribosomal protein (L36, L19, L10, L25) (*RS_12165*, *RS_06670*, *RS_18765*, and *RS_18825*) and 30S ribosomal proteins (S8, S10, S21) (*RS_18645*, *RS_18720*, and *RS_21170*) were also found to be downregulated **Figure 6(B)**.

#### qRT-PCR validation

To confirm RNA-seq data’s reliability, the expression level of eight genes, which included four upregulated genes (*RS_01475*, *RS_02860*, *RS_06840*, and *RS_19500*) and four downregulated genes (*RS_02865*, *RS_19815*, *RS_19820*, and *RS_15130*) was examined using qRT-PCR, shown in **Supplementary Figure 4**. For all the genes, the same expression trend was detected for both qRT-PCR and RNA-seq analyses. Up-regulation of genes that constitutes *ars* operon in qRT-PCR **Figure 4(E)** further confirmed the reproducibility and reliability of RNA-seq data.

## Discussion

Heavy metal contamination in the environment is a serious threat to human health. Metals such as arsenic, cadmium, chromium, lead, and mercury are commonly occurring metals and rank among the priority metals of public health risk. Among these metals, arsenic contamination in drinking water and food crops, especially in rice, is a growing concern, particularly in countries like India, Bangladesh, and China (Mondal et al., 2020) (Mondal and Polya, 2008) (Nickson et al., 2000). Unlike in any other crops, water embedded conditions in the paddy fields make the environment suitable for the accumulation of inorganic forms of arsenic, specially arsenite As(III) that is highly toxic to the cells (Mitra et al., 2017) (Takahashi et al., 2004). Studies have shown the transport and accumulation of arsenic to the different parts of the rice plants, including roots, shoots, straw, and grains (Suriyagoda et al., 2018) (Ma et al., 2008) (Wu et al., 2011). Rice is an efficient accumulator of arsenic compared to the other crops due to arsenite’s mobilization and sequestration of arsenite through iron oxides under anaerobic conditions (Yamaguchi et al., 2011). Rice is the staple food for the Indian population, and human exposure to arsenic through rice intake is causing serious health issues. In the present study, we have elucidated the arsenic resistance mechanisms of *Xanthomonas oryzae* pv. *oryzae* (*Xoo*), which is a serious and highly-specific rice pathogen. The presence of metals in the environment can have influence on the survival of pathogens as well as plants. In this study, we highlighted the interplay between plant, pathogen, and environmental interactions. Our study highlights the important and specific role of unique micro-environmental conditions, like in submerged rice cultivation, can lead to high arsenic stress to both host and microbes associated with it. In this context, acquired resistance against high arsenic stress by acquiring novel genomic cassettes in an emerging lineage indicates ongoing selection pressure on this host-specific pathogen. The absence of the cassette in other lineages of *Xoo* and at the same time in a lineage that is predominant to the Asian continent reiterates unique selection pressure on the *Xoo* population.

We observed a horizontally acquired cassette in the *Xoo* IXO1088 encoding genes involved in the arsenic metabolism. The cassette includes an *ars* operon regulated by an arsR family regulator and is induced in the presence of both arsenate and arsenite. The cassette provides high-level resistance to *Xoo* IXO1088 against both arsenate and arsenite. Further spread of this cassette in a particular lineage suggests the recent evolution and success of strains belonging to this particular lineage to adapt against arsenic toxicity. To better understand the adaptive response of *Xoo* population to arsenic exposure and toxicity coping mechanisms, we performed a transcriptome-sequencing study. RNA-Seq is an efficient high-throughput sequencing method to study bacterial transcriptomes and explore various pathways involved in arsenic resistance mechanisms. Our transcriptome-based study of *Xoo* under arsenic stress highlights major pathways differentially expressed in *Xoo* IXO1088 in response to arsenate and arsenite.

Our RNA-Seq analysis revealed that the gene expression patterns in *Xoo* IXO1088 were different in both arsenate and arsenite treated samples. Commonly upregulated genes under arsenate and arsenite treatment include upregulation of *ars* operon that encode proteins involved in arsenic detoxification. Other than this MarR and Abr family transcriptional regulators and proteins involved in iron acquisition, transport, and storage such as siderophores and bacterioferritin were also upregulated, consistent with the previous studies (Zhang et al., 2016). In arsenic conditions, more iron is recruited to alleviate stress. In flooded rice fields, iron is in ferric (Fe^3+^) (insoluble oxidized) form, which is converted to ferrous (Fe^2+^) form that sequesters arsenic (Takahashi et al., 2004). Production of siderophores is the common strategy for iron acquisition under Fe-limiting conditions. Some metals also possess affinity towards siderophores. The complex formation of siderophores with metals reduces complexation with iron, thus decreasing the concentration of soluble iron. Iron deficiency subsequently leads to the production of siderophores (Schalk et al., 2011).

A relatively large number of genes were expressed in response to arsenite compared to arsenate treatment. Many differentially expressed genes specific to arsenite encoding proteins involved in the oxidative response, protein damage, ion transport, recombination and energy generation, cell motility and signal transduction were found to be upregulated. Arsenic exposure induces reactive oxygen species (ROS) and nitric oxide (NO) production inside the cell, and to reduce the oxidative damage, bacteria induce various antioxidant enzymes (Andres and Bertin, 2016) (Maity et al., 2012). We observed upregulation of superoxide dismutase, catalase, organic hydroperoxide resistance protein, alkyl hydroperoxide reductase subunit F (aphF), oxidative stress transcriptional regulator that participate in the control of elevated ROS levels. Another adaptive response to arsenite treatment in *Xoo* IXO1088 includes the upregulation of various heat shock proteins (HSPs). HSPs help in survive under stress conditions by facilitating the proper folding of newly or already translated proteins and preventing the accumulation of aggregated proteins. Upregulation of chaperons has also been observed in response to other metals (Tamás et al., 2018) (Elran et al., 2014). Chaperons can contribute by converting metal-induced misfolded proteins to active native proteins, thus enhancing cellular tolerance to heavy metals. Various transporters, including ATP-binding cassette proteins, MFS (major facilitator superfamily), ferrous ion transport protein were also upregulated. Additionally, chemotaxis is one of the most important features of bacteria that regulate their movement towards or away from specific substances, either attractants or repellents. The transcriptome data also showed upregulation of four DEGs CheW, CheA, motD, and methyl-accepting chemotaxis protein, suggesting role of chemotaxis response towards arsenite. Upregulation of genes involved in carbohydrate metabolism pathways was also in agreement with the previous studies (Cleiss-Arnold et al., 2010) (Shah and Damare, 2020).

The majority of the genes encoding proteins translational machinery (ribosomal proteins) and proteins involved in amino acid metabolism, were downregulated in response to arsenite. This finding is consistent with the previous studies showing a decline in the rate of protein synthesis in response to arsenite As(III) exposure (Wang and Crowley, 2005) (Shah and Damare, 2020) (Henne et al., 2009). In addition, genes involved in oxidative phosphorylation pathways such as ATP synthase subunits A, C, delta and beta, cytochrome ubiquinol oxidase subunits, and genes involved in intracellular protein transport including *secD*, *secF*, *secG*, and *secE* were also downregulated in response to arsenite treatment. Multi-drug RND-efflux transporter subunits and potassium ATPase subunit F and A were downregulated in both the conditions. The downregulation of these genes indicate switch from aerobic to anaerobic respiration to promote energy conservation following arsenic exposure.

In conclusion, we have identified an evolved cassette in a lineage of *Xoo* population that confers arsenic resistance. The arsenic-rich environment has possibly led to the acquisition of such cassette in *Xoo* population and can provide selective/adaptive advantage to the highly virulent strains. Using RNA-Seq technology, we have identified the adaptive and detoxification pathways adapted by *Xoo* to evade arsenic toxicity. The study highlights the evolution of pathogens to adapt to unique microenvironment during course of evolution and diversification into different lineages. Altogether, the study could be useful for further studies to understand molecular mechanisms under arsenic stress and manifest the need to tackle arsenic contamination in rice fields.

## Materials and methods

### Bacterial growth and Arsenic sensitivity assay

*Xanthomonas oryzae* pv. *oryzae* strains IXO1088 and BXO512 were used in the present study to represent arsenic resistance and sensitive strains, respectively. List of strains, vectors and primers used in the study is given in **Supplementary Table 3**. The complete genome sequence of IXO1088 is available at NCBI with accession number CP040687, and the complete genome of BXO512 was sequenced using Nanopore sequencing technology as described in the previous study (Kaur et al., 2019). The BXO512 genome was submitted to NCBI with accession number CP065228. For sensitivity assay, strains were grown in nutrient broth (NB) media at 28 °C at 180 rpm. Cultures were diluted to an optical density of 1.0 at OD_600_. One percent of the cultures was added to the fresh 10 ml NB aliquots containing increasing concentrations of sodium arsenate (Na_2_HAsO_4_.7H_2_O) or sodium arsenite (NaAsO_2_) (purchased from Himedia Co.) added at zero time point. Then, cultures were incubated for 48hrs, and after that, growth was measured at OD_600_. Cultures without arsenate/arsenite treatment were used as control. For growth curve assay, cultures were treated with given concentrations of arsenate and arsenite and incubated for 78 hrs. Samples were taken at regular intervals (6hrs), and growth was measured as OD_600_.

### Sequence characterization and distribution of a novel*ars* cassette in*Xoo* population

Putative promoter sequences were identified using the BPROM software (http://www.softberry.com/berry.phtml?topic=bprom&group=programs&subgroup=gfindb). Protein BLAST (Basic local alignment search tool) was used to find out the homologs of ArsR protein. Homolog proteins obtained from different species were then used for multiple sequence alignment (MSA) using ClustalW (https://www.genome.jp/tools-bin/clustalw). Phylogenetic analysis was performed using MEGA6 software by the neighbor-joining method (Tamura et al., 2013). A core genome tree of all *Xoo* genomes available at NCBI was constructed using PhyML (Guindon et al., 2010). Briefly, Core genome alignment was obtained using Roary v3.11.2 (Page et al., 2015) and converted to phylip format using SeaView v4.4.2-1 (Gouy et al., 2010). Then, PhyML was used to obtain a Newick tree file, which was visualized using iTOL (Letunic and Bork, 2016). The presence of *ars* cassette in other *Xoo* population was scanned by taking cassette sequence as a query and performing nucleotide BLAST against all the *Xoo* genomes available in the NCBI.

### RNA sequencing and data analysis

The sublethal concentrations of arsenate and arsenite were selected by growing cultures in the different concentrations and measuring growth at regular time intervals. Strains were grown in nutrient broth media at 28□ C at 180 rpm to an OD_600_ of 1.0. One percent of the aliquot was added to the fresh media and grown-up to OD_600_=0.4; then cultures were treated with different concentrations of arsenate (0.5mM and 1mM) or arsenite (0.6mM) for 30 minutes. Culture without treatment was used as control. The cells were harvested by centrifugation at 6,000g for 5min for RNA isolation. Total RNA was isolated using Direct-zol RNA MiniPrep kit (Zymo Research Corporation, Orange, CA, USA), and samples were shipped to Agrigenome Labs Pvt. Ltd. (India). Briefly, the purity of isolated RNA was determined using the NanoDrop(Thermo Scientific, Wilmington, DE, USA) and quantified using Qubit(Invitrogen, Carlsbad, CA, USA). RNA integrity was checked using Agilent 2200 Tape Station system (Agilent Technologies, Netherlands). The rRNA was removed using ‘si-tools Pan-prokaryote ribopool probes,’ and further library preparation was done using TruSeq Stranded RNA library prep kit. Prepared libraries were sequenced using Illumina HiSeq x10 platform, and 150-bp paired-end reads were generated. Clean reads that passed the quality filter were mapped to the reference genome (*X. oryzae* pv. *oryzae* IXO1088 accession number CP040687) using EDGE pro (version-1.3.1). Differential gene expression analysis of treated samples with respect to control was performed using DESeq program. Differentially expressed genes with adjusted *p-*value <0.05 and log_2_fold change value ≥1or ≤−1 were used for further analysis. Common and unique DEGs were visualized by Venn diagram and Volcano plots using the R software package. Heat maps were generated using log_2_fold change values of treated versus control samples using GENE-E software. DEGs with log_2_fold change value ≥1or ≤−1 were classified into different COG classes using the eggNOG tool (Huerta-Cepas et al., 2017).

### Validation of RNA-Seq data by qPCR

To check the induction of *ars* cassette and to validate RNA-Seq data, quantitative real-time PCR (qPCR) experiments were performed. The primers used are listed in **Supplementary Table 3**. The quantitative real-time PCR assay was performed with SuperScript III Platinum SYBR Green One-Step qRT-PCR kit (ThermoScientific, Wilmington, DE, USA). Three technical replicates were included for each sample, and reactions were set up according to the manufacturer’s instructions. The amplification conditions were: cDNA synthesis 50°C for 45 min, initial denaturation at 95 °C for 5 min, 40 cycles of denaturation at 95 °C for 15s followed by annealing at 60 °C for 30 s and extension at 40 °C for 30 s. ftsZ was used as an endogenous control. Each gene’s relative expression in the treated and untreated sample was expressed as a fold change calculated using the 2^−ΔΔct^ method. Three biological replicates were included for each condition.

### Cloning of *ars* cassette

*Xoo* strain IXO1088 genomic DNA was isolated from overnight grown culture using Zymo kit. The complete *ars* gene cassette was amplified using primers listed in **Supplementary Table 3**. PCR conditions were as follows: initial denaturation 95 □C for 30s; 35 cycles of denaturation 95 □C for 10s; annealing 60 □C for 30s; extension 72 □C for 3min; and final extension 72 □C for 10min. The amplified fragment was gel extracted using the QIAquick Gel Extraction Kit, given kinase treatment, purified, and ligated into the pUFR034 vector. The vector was linearized using inverse M13 primers. The ligation mix was transformed to *E. coli* Top10 cells. The recombinant plasmid was verified using M13 forward and reverse universal primers and insert specific primers. For transformation of *E. coli* cells, ligation mix was added to the 100μl of competent cells and incubated on ice for 15 min. Then, competent cells were subjected to heat shock at 42 □C for 60 seconds. Cells were immediately shifted to the ice for 2 min. Then 1ml of LB media was added, and cells were incubated at 37 □C at 200 rpm for 1hr. After incubation, cells were plated on LB agar plates containing kanamycin (50μg/ml). Plates were incubated at 37 □C for 12 hrs for the growth of recombinants.

### Preparation of electrocompetent cells and transformation of *X. oryzae* pv. *oryzae*

For the preparation of electrocompetent cells, *Xoo* strain BXO512 was inoculated in 6ml nutrient broth (NB) media and cultured till late exponential phase (OD = 0.8-1.0) at 28□ C with 180rpm shaking. The culture was then equally divided into four centrifuge tubes, and cells were harvested by centrifugation at 16,000g for 2 min at room temperature. The pellets in each tube were then washed with 1ml of 300mM sucrose. The washing step was repeated three times at the same speed. Finally, cells in four tubes were resuspended with a total 100μl of 300mM sucrose. 100μl aliquot of the competent cells was used for one electroporation experiment.

For electro-transformation, 100μl of electrocompetent cells was mixed with 10μl (500ng-1μg) of plasmid DNA and transferred to 2mm ice-cold electroporation cuvettes (Gene Pulser Cuvette (BioRad) and tapped gently to avoid air bubbles. Cells without plasmid were used as a negative control. Further, electrocompetent cells were subjected to electroporation at 2.5kV, 25μF, and 200 Ω using the Gene Pulsar Xcell electroporation system (BioRad Laboratories, Inc.). After the electric shock, immediately, 1ml of NB broth was added to the cuvette, mixed properly, and transferred to the 5ml micro-centrifuge tubes. The electroporated cells were incubated at 28 □C at 180 rpm for 6 hrs. After recovery, cells were plated on nutrient agar plates containing kanamycin antibiotic (15μg/ml) and incubated at 28 □C until colonies appeared (usually 72hrs).

### Data deposition

The genome sequence of strain BXO512 is submitted to NCBI with accession number CP065228.

## Abbreviations

Xoo: *Xanthomonas oryzae* pv. *oryzae;*
As: arsenic
Hsps: Heat shock proteins
DEGs: Differentially expressed genes
COG: cluster of orthologous groups
ROS: reactive oxygen species
qPCR: quantitative real-time PCR
MSA: multiple sequence alignment

## Authors Contribution

AK performed all the experiments and analyzed the results with the help of RR. RR performed the qRT-PCR experiment to validate RNA-Seq data. TS helped in analyzing RNA-Seq data. AK drafted the manuscript with inputs from RR, TS, and PBP. PBP and AK have conceived and participated in designing the study. PBP applied for the funding. All authors have read and approved the manuscript.

## Acknowledgements

This work was supported by a CSIR project entitled “High throughput and integrative genomic approaches to understand adaptation of probiotic and pathogenic bacterium” (OLP148). AK is supported by DST-INSPIRE fellowship. RR and TS are supported by CSIR fellowship.

## Supplementary material

**Supplementary Figure 1:** Distribution of ars cassette in different states in **(A)** India **(B)** across Asian countries. The black dots represent the presence of *ars* cassette.

**Supplementary Figure 2:** Growth curve assay: Growths curves of *X. oryzae* IXO1088 strain in the presence of different concentrations of **(A)** sodium arsenate or **(B)** sodium arsenite. The values represent the average of replicate experiments with standard deviations from three independent cultures.

**Supplementary Figure 3:** Volcano plots of differentially expressed genes between control and arsenate/arsenite treated groups. The red dots represent significantly (adjusted *p-*value < 0.05) upregulated (log_2_fold change ≥1) and downregulated (log_2_fold change ≤−1) genes, blue dots represent genes with adjusted *p-*value < 0.05 but that do not meet log_2_ fold change cut-off, green and black dots means insignificant gene expression with *p-*value > 0.05.

**Supplementary Figure 4:** Validation of RNA-Seq data using qPCR. Eight commonly up and downregulated genes under both conditions were selected. The light grey bars represent the log2 fold change of RNA-Seq data. The black bars represent the mean log2 fold change of qPCR data obtained from two biological replicates with error bars showing standard deviation.

**Supplementary Table 1:** Showing GC% content of genes present in the *ars* cassette.

**Supplementary Table 2:** List of DEGs (differentially expressed genes) upregulated and downregulated with log_2_fold change ≥1 and log_2_fold change ≤−1.

**Supplementary Table 3:** List of primers used in this study.

