## Supplementary figures and images for "Discerning role of a functional arsenic resistance cassette in evolution and adaptation of a rice pathogen"

### Supplementary Figure 1

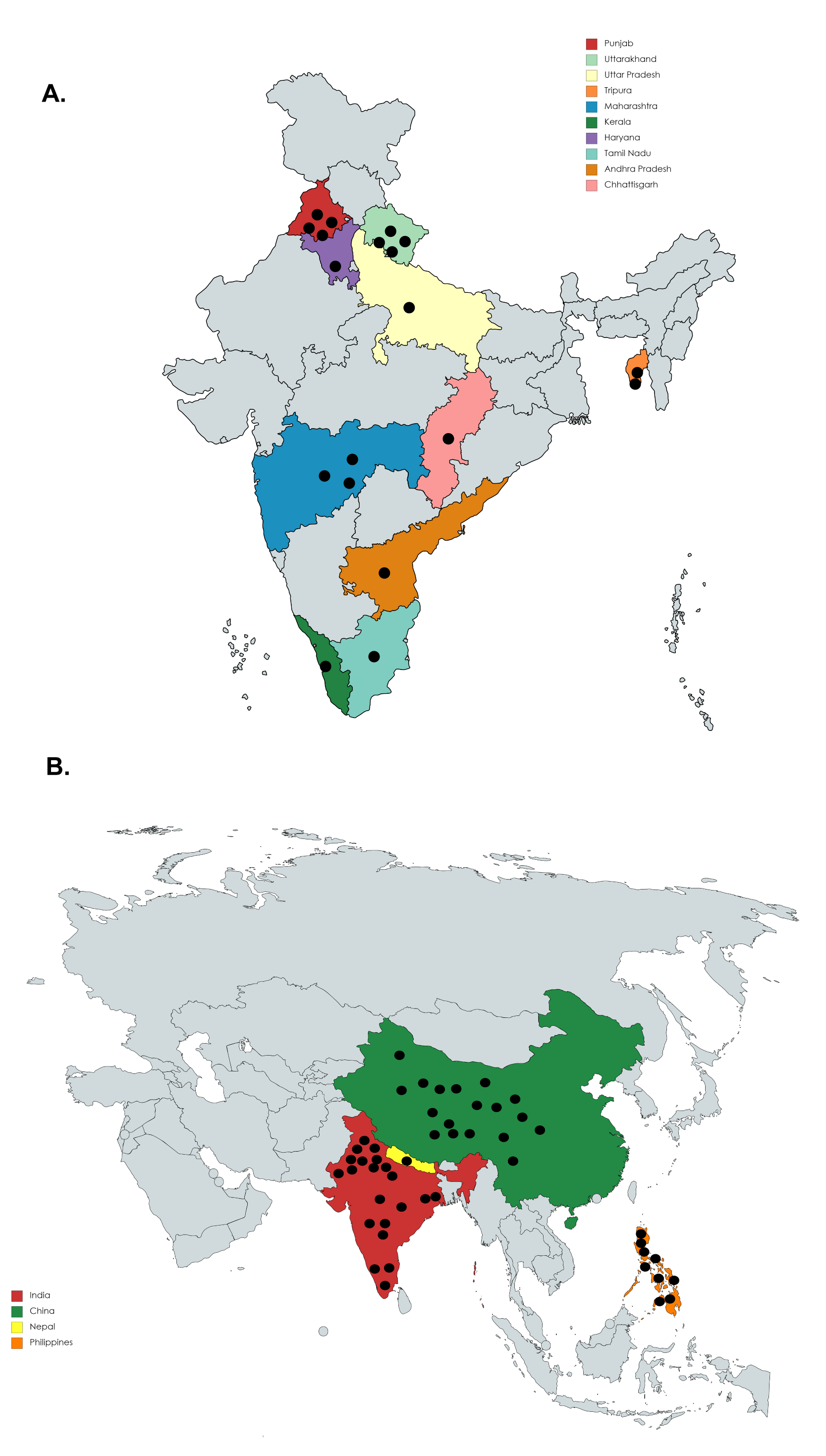

### Supplementary Figure 2

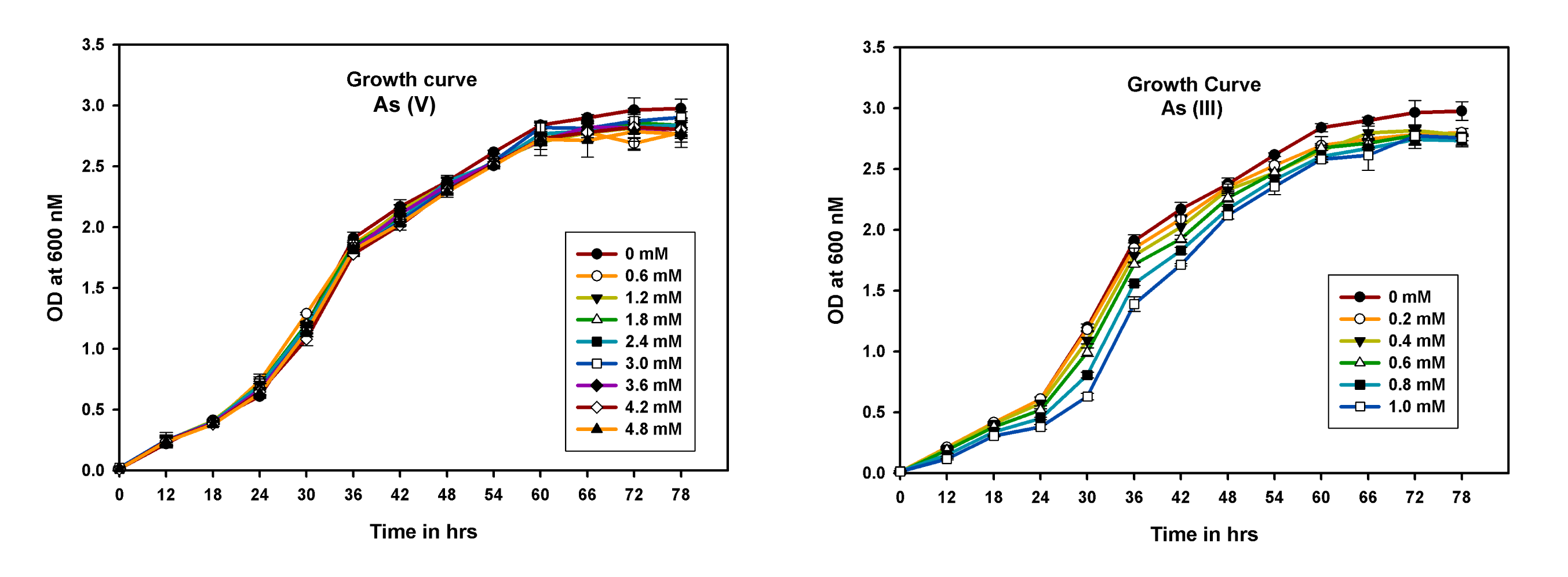

### Supplementary Figure 3

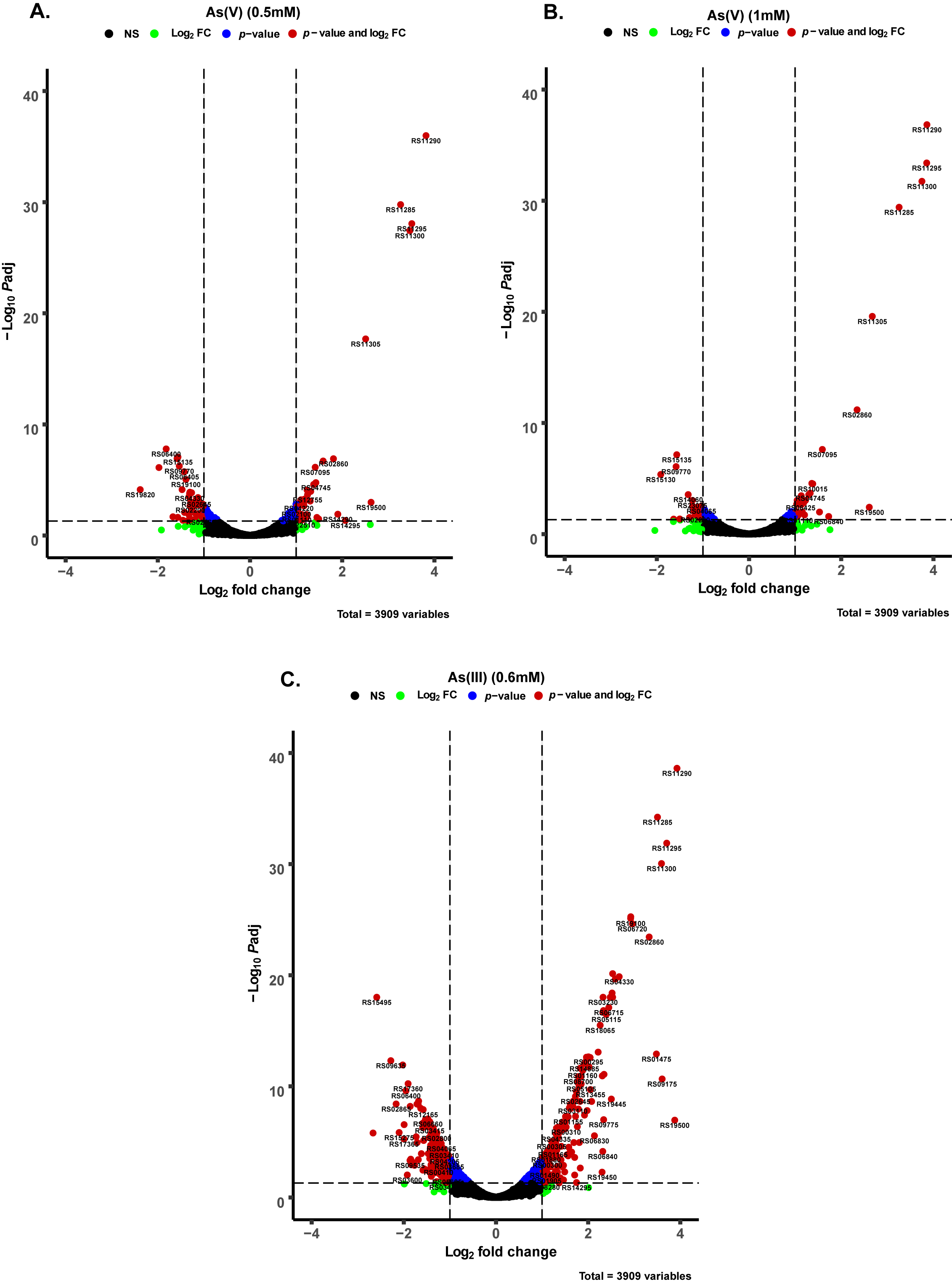

### Supplementary Figure 4

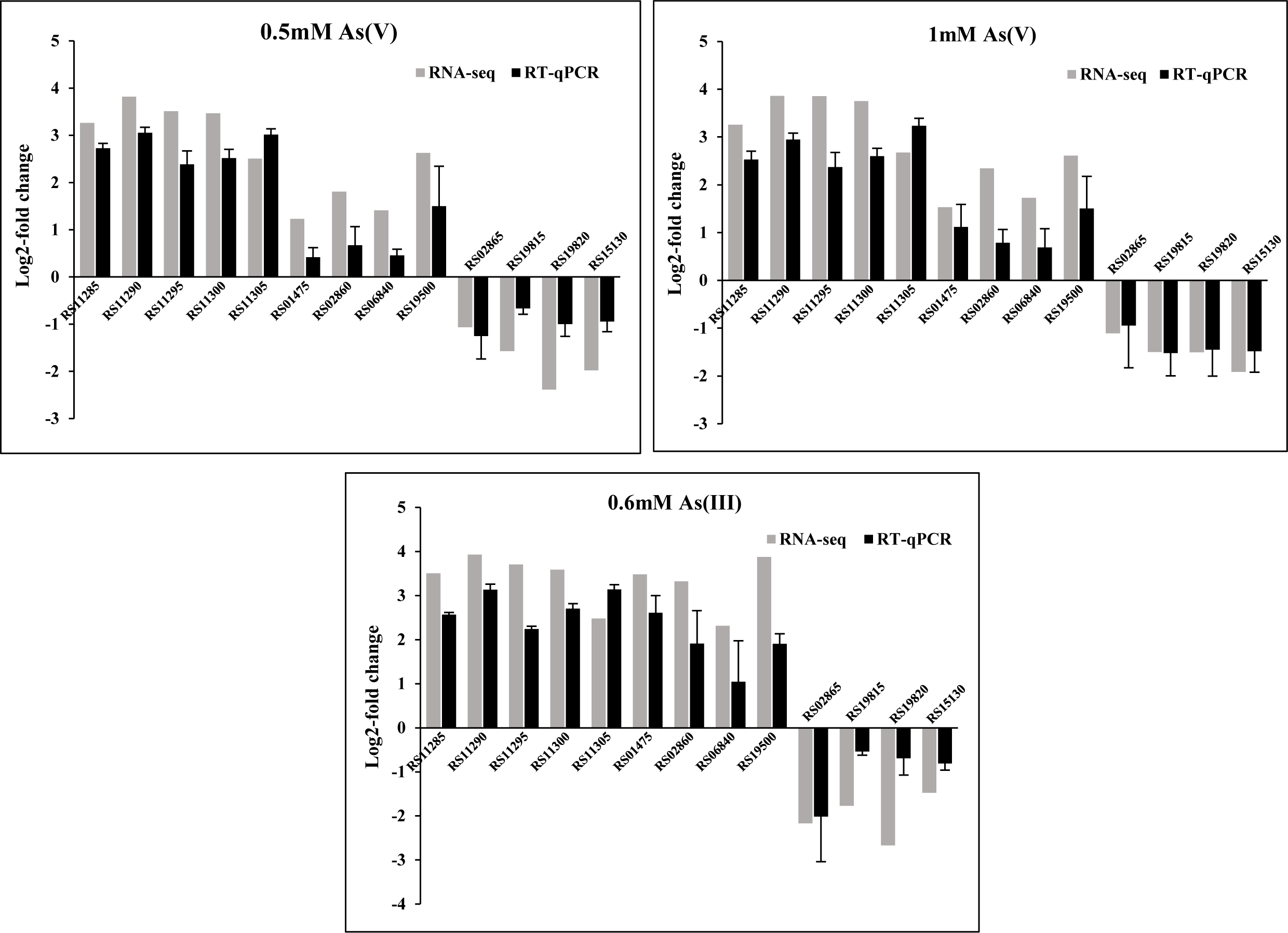
